# Multi-omics approach reveals dysregulation of protein phosphorylation correlated with lipid metabolism in mouse fatty liver

**DOI:** 10.1101/2022.02.16.480672

**Authors:** Sora Q. Kim, Rodrigo Mohallem, Jackeline Franco, Kimberly K. Buhman, Kee-Hong Kim, Uma K. Aryal

## Abstract

Obesity caused by overnutrition is a major risk factor for non-alcoholic fatty liver disease (NAFLD). Several lipid intermediates such as fatty acids, glycerophospholipids and sphingolipids are implicated in NAFLD, but detailed characterization of lipids and their functional links to proteome and phosphoproteome remain to be elucidated. To characterize this complex molecular relationship, we used multi-omics approach by conducting comparative proteomic, phoshoproteomic and lipidomic analyses of high fat (HFD) and low fat (LFD) diet fed mice livers. We quantified 2447 proteins and 1339 phosphoproteins containing 1650 class I phosphosites (with localization probability > 0.75), of which 669 phosphosites were significantly different between HFD and LFD mice livers. We detected alterations of proteins associated with cellular metabolic processes such as small molecule catabolic process, monocarboxylic acid, long- and medium-chain fatty acid, and ketone body metabolic processes, and peroxisome organization. We observed significant downregulation of protein phosphorylation in HFD fed mice liver in general. Untargeted lipidomics identified upregulation of triacylglycerols, glycerolipids and ether glycerophosphocholines and downregulation of glycerophospholipids such as lysoglycerophospholipids, as well as ceramides and acylcarnitines. Analysis of differentially regulated phosphosites revealed phosphorylation dependent deregulation of insulin signaling as well as lipogenic and lipolytic pathways during HFD induced obesity. Thus, this study reveals a molecular connection between decreased protein phosphorylation and lipolysis, as well as lipid-mediated signaling in diet-induced obesity.

## 1. Introduction

Obesity is a prevalent health concern worldwide and it is accompanied by a plethora of comorbidities. Among them, non-alcoholic fatty liver disease (NAFLD) is now recognized as the most common form of liver disease affecting one quarter of the global population, which is similar rate of prevalence as obesity [1]. NAFLD is characterized by fat accumulation in the liver that is not caused by alcohol consumption, and is associated with different factors, including increase in dietary fat released from adipocytes via lipolysis, de novo hepatic lipogenesis, circulating free fatty acid (FFA) and decrease in fatty acid oxidation [2,3]. NAFLD poses a significant health burden because even the earliest and most common type of NAFLD, simple steatosis, is shown to be not as prognostically benign as it has been thought for a long time. As an illustration, recent large-scale cohort study of 10,568 patients with NAFLD in Sweden found that patients with simple steatosis showed a significantly elevated risk of extrahepatic cancer, hepatocarcinoma, cardiovascular disease and cirrhosis [4]. Therefore, it is critical to understand the molecular signatures involved in the onset of steatosis as failure of proper liver functions will promote pathogenesis of metabolic complications in turn [5].

HFD feeding is advantageous for the establishment of fatty livers in mice as it generates less pronounced inflammation and rarer fibrosis after extended period of intervention than in case of methionine-choline deficient diet [6,7]. Moreover, the model establishes pathologic phenotypes resembling human disease as it is accompanied by obesity, insulin resistance and hyperlipidemia [6,7]. Therefore, the diet-induced obesity (DIO) is typically associated with complex and intertwined metabolic abnormalities that primarily entail the involvement of a multitude of proteins, including their differential expression, post-translational modifications, and protein-protein interactions [8–10]. Like proteins, lipids also have extensive biological roles as signaling molecules, energy reserves and structural components of membranes. Gaining information about regulation/dysregulation of each class of lipids in response to DIO and integrating these results with proteomics and phosphoproteomics is important for our comprehensive understanding of the etiology of fatty liver. In the last three decades, significant progress has been made in our understanding of the wide-ranging changes in proteins, RNA and metabolites caused by obesity and overnutrition [8,11–14]. However, most of the omics studies on obesity and insulin resistance thus far have focused on one area (either proteomics or lipidomics), and only a few have focused on multi-omics approaches. In particular, studies integrating global proteomics, phosphoproteomics and lipidomics of liver under the context of DIO are limited. Such integrated analyses, however, can be a powerful strategy to depict the changes in cellular functions controlled by highly connected protein and lipid molecules. This knowledge could be particularly useful for understanding the biology of obesity-related pathologies, and for the development of new treatment strategies.

In the present study, we aimed to explore the changes in liver proteins and lipids and their functional role and relationship in the development of DIO at systems level by performing an integrative multi-omics analysis and correlating identified modules with DIO. Our comprehensive analysis revealed altered expression of many key metabolic enzymes, membrane transporters and kinases that are directly or indirectly involved in lipogenesis and lipolysis, reflecting metabolic imbalance of lipid homeostasis during the development of HFD induced obesity and insulin resistance. Particularly, we observed loss of phosphorylation on acetyl-coenzyme A synthetase (Acss2), Acetyl-CoA carboxylase (Acaca, Acacb), ATP citrate lyase (Acly), fatty acid synthase (Fasn), sugar transporter 2 (Glut2) and many other key enzymes of lipid metabolism and energy generation. Lipidomic analysis revealed increased accumulation of triacylglycerols, glycerolipids and ether glycerophosphocholines and decreased accumulation of glycerophospholipids as well as ceramides and acylcarnitines. Free fatty acids and ether glycerophosphocholines were negatively correlated with most of the significantly disregulated phosphosites while phosphatidylcholine and lysophosphatidylcholine were positively correlated. This suggest the importance of the lipid environment for the membrane transporters and kinases functionality, and highlights the relevance of mechanistic studies in this domain.

## 2. Materials and Methods

### 2.1. Mouse husbandry and diets

All procedures involving animals were performed in accordance with the National Institute of Health Guide for the Care and Use of Laboratory Animals and were reviewed and approved by the Purdue Animal Care and Use Committee (protocol number 1111000154). C57BL/6 male mice were kept in a humidity and temperature-controlled facility in a 12:12 h dark/light cycle with *ad libitum* access to food and water. From weaning to 5 weeks of age, mice were fed a chow diet consisting of 62.1% of calories from carbohydrate (starch), 24.7% from protein, and 13.2% from fat (PicoLab 5053, Lab Diets, Richmond, IN, United States). At 5 weeks of age, mice were randomly assigned to one of the two diets for additional 12 weeks: low-fat diet (LFD, 10% calories from fat, D12450J) or high-fat diet (HFD, 60% calories from fat, D12492) (Research Diets, Inc.; New Brunswick, NJ, United States). Body weight was recorded weekly. Fasting blood glucose was measured using the OneTouch glucometer (LifeScan, Milpitas, CA). After 12 weeks, mice were fasted for two hours and euthanized by CO_2_ followed by cervical dislocation and livers were collected and stored at −80 °C for further analysis. Fat pad weight was measured at the time of euthanasia.

### 2.2. Liver tissue preparation for proteomics

Livers from six lean mice and six DIO mice were homogenized in 100 mM ammonium bicarbonate, supplemented with protease and phosphatase inhibitors, in a Precellys Evolution using CK14 soft tissue homogenizer tubes (Bertin Technologies SAS) for 3 × 20s bursts at 6200 rpm. Protein concentration of each homogenate was determined by bicinchoninic acid assay kit (Thermo Fisher Scientific). An aliquot of each sample containing 500 ug of total protein was precipitated using ice-cold acetone at −20 °C overnight. After acetone removal, protein pellets were reduced with 10mM dithiothreitol in 8M urea, and alkylated using iodoethanol in ACN (2% iodoethanol, 0.5% triethylphosphine, 97.5% acetonitile). Proteins were digested with mass spec grade Trypsin/LysC mix (Promega) at a 1:50 (w/w) enzyme-to-substrate ratio, using a barocycler (Pressure BioScience Inc.) at 50°C with 60 cycles of 20kpsi for 50 seconds and 1 atmospheric pressure (1 ATM) for 10 seconds. Samples were cleaned using Pierce Peptide Desalting Spin Columns (Thermo Fisher Scientific). For phosphopeptide enrichment, PolyMac phosphopeptide enrichment spin-tips (Tymora Analytical) were used following manufacturer’s recommendations.

### 2.3. Mass spectrometry analysis of liver proteome

Samples were analyzed by reverse-phase LC-ESI-MS/MS system using the Dionex UltiMate 3000 RSLC nano System coupled to the Orbitrap Fusion Lumos Mass Spectrometer (Thermo Fisher Scientific, Waltham, MA) as described previously [14,15]. Briefly, peptide separation was accomplished using a trap column (300 μm ID × 5 mm) packed with 5 μm 100 Å PepMap C18 medium, and then using a reverse phase analytical column (50-cm long × 75 μm ID) packed with 2 μm 100 Å PepMap C18 silica (Thermo Fisher Scientific, Waltham, MA). The column was maintained at 50 °C, mobile phase solvent A was 0.1% formic acid (FA) in water and solvent B was 0.1% FA in 80% acetonitrile (ACN). Peptides were separated using a 160-min LC gradient at a flow rate of 150 nL/min. The mass spectrometer was operated in positive ion and standard data-dependent acquisition mode with Advanced Peak Detection function activated. The fragmentation of precursor ion was accomplished by higher energy collision dissociation at a normalized collision energy setting of 30%. The resolution of Orbitrap mass analyzer was set to 120,000 and 15,000 at 200 m/z for MS1 and MS2, respectively, with maximum injection time of 50 ms for MS1 and 20 ms for MS2. The dynamic exclusion was set at 60s to avoid repeated scanning of identical peptides and charge state was set at 2-7 with 2 as a default charge. The full scan MS1 spectra were collected in the mass range of 375-1,500 m/z and MS2 in 300-1250 m/z. The spray voltage was set at 2 and Automatic Gain Control (AGC) target of 4e5 for MS1 and 5e4 for MS2, respectively.

### 2.4. Data analysis

LC–MS/MS data were processed with MaxQuant software (Ver 1.6.17.0). Raw spectra were searched against the mouse UniProt FA *Mus musculus* protein database (June 2021). Six amino acids were set as the minimum length required in the database search. The search was performed with a precursor mass tolerance of 10 ppm and MS/MS fragment ions tolerance of 20 ppm. Trypsin and LysC were set as specific enzymes, with up to two missed cleavages allowed. Oxidized methionine, and for the phosphoproteomics, phospho STY were defined as a variable modification, and iodoethanol of cysteine was defined as a fixed modification. The “unique plus razor peptides” were used for peptide quantitation and the false discovery rate of peptides spectral match and protein identification was set at 1%. “Lable free quantitation” (LFQ) and “Match between runs” were enabled. Subsequent bioinformatics analysis was performed with Perseus (ver 1.6.7.0). Proteins labeled “only identified by site”, “reverse”, or “contaminants” were removed from the analysis. Proteins were filtered for at least five valid values among six biological replicates (70%) in either of conditions. Missing values were replaced by values derived from a normal distribution. Significantly up-or-downregulated proteins between the two groups were determined by Student’s t-test with Permutation-based FDR (q-value < 0.05, log(FC) > 0.38). For phosphoproteomics, only class one phosphosites (localization probability > 0.75) were considered for downstream analysis. Functional annotations and enrichment analysis were performed using Metascape.

### 2.5. Lipid extraction and measurement

Lipids were extracted using the Bligh & Dyer extraction method [16]. Briefly, 200 μL liver homogenates with equal amount of protein were transferred to a new tube and mixed with 550 μL methanol and 250 μL of chloroform. The solution was vortexed briefly and incubated at 4 °C for 15 min, 250 μL of ultrapure water and 250 μL of chloroform were added and the sample was centrifuged for 10 min at 16,000 x g, forming a 2-phase solution. Bottom organic phase was transferred to a new tube and dried and samples were stored at −80 °C until analysis. Lipid extracts were analyzed in an Agilent 1290 Infinity II UPLC, coupled to an Agilent 6545 quadrupole time-of-flight tandem mass spectrometer. Samples were resuspended in 30 μL of a mixture of ACN: methanol: water (3 :5 :2 v/v) and 8 μL were loaded to a Waters ACQUITY UPLC® BEH C18 1.7 μm column with controlled temperature of 45°C. The binary pump used 10mM ammonium acetate in water with 0.1% formic acid for mobile phase A and 10mM ammonium acetate in a 50% isopropyl alcohol: 49.9% acetonitrile: 0.1% formic acid for mobile phase B at a flow rate of 0.4ml/min. The liquid chromatography gradient was of 35% B at 0 minutes, 80% at 5 minutes and ramped up to 100% B at 10 minutes, with a 5-minute hold, then returned to 35% in 2 minutes and a 4-minute hold. The mass analyzer was set with a ESI capillary voltage of 35000Vcap, a sheath gas temperature of 320°C and flow of 8L/min, a nebulizer gas pressure of 35psig, a fragmentor of 135Vs and skimmer of 35 V. Mass spectrums were collected in profile mode with a range from 100 to 1200 m/z at a scan rate of 5 spectra/s with 200 min/spectrum for MS1 and 3 spectras/s with 333.3 min/spectrum for MS2. Raw data were processed using MS-dial 4.7 [17] with the MSP spectral kit of 13,303 unique compounds in positive mode.

A second lipid extract obtained as described above was used for fatty acid methylations (FAMEs) to analyze free fatty acid content by GC/MS. Before derivatization, samples were spiked with 1ug of C17:0 as internal standard. The samples were derivatized with 500uL of 14% boron trifluoride solution (Sigma-Aldrich # B1252) and reacted for 30 minutes at 60°C followed by the addition of 500uL of water and 500uL of hexane. After mixing, 0.2g of anhydrous sodium sulfate was added to the sample and let it sit. The hexane layer was collected and dried. GC/MS derivatized samples were resuspended in 100uL of 100% hexane in a Thermo Fisher Triplus RSH auto sampler and Trace 1310 gas chromatography (GC) system coupled to a Thermo Fisher TSQ 8000 mass spectrometer (MS) (Thermo Fisher Scientific, Waltham, MA) with a column Agilent Select FAME GC column (50 m x 0.25 mm, film thickness 0.2 um) (Agilent Tech-nologies, Santa Clara, CA). The GC carrier gas was helium with a linear flow rate of 1.5 ml/min. The GC temperature gradient started at 70°C at 0 minute and ramped to 310°C at 7°C/min and hold for 1 minute for a total run time of 38.28 minutes. The GC inlet was set to 250°C and samples were injected in split mode with a ratio of 10 and a flow of 15 ml/min. The MS transfer line was set to 250°C and the MS ion source was set to 250°C. MS data were collected in selected ion monitoring (SIM) mode. Raw data were analyzed with Thermo Fisher Chromeleon (Version 7.2.9) software and a standard mixture of 37 FAME (Sigma-Aldrich Corp., St. Louis, MO) was used to confirm spectra and column retention times.

## 3. Results

### 3.1. Mice fed a chronic high-fat diet develop obesity

Male C57BL/6J mice were fed either a LFD or a HFD for 12 weeks starting at 5 weeks of age. After only 1 week, body weight of HFD fed group started to be significantly greater than that of LFD fed group (*P* < 0.05) and after 12 weeks of diets, we confirmed that HFD induced mice obesity (weight gain of 238%). HFD also increased adipose tissue fat deposition and elevated fasting blood glucose concentrations, which indicate defects in blood glucose control (Figure 1B, C).

**Figure 1.**
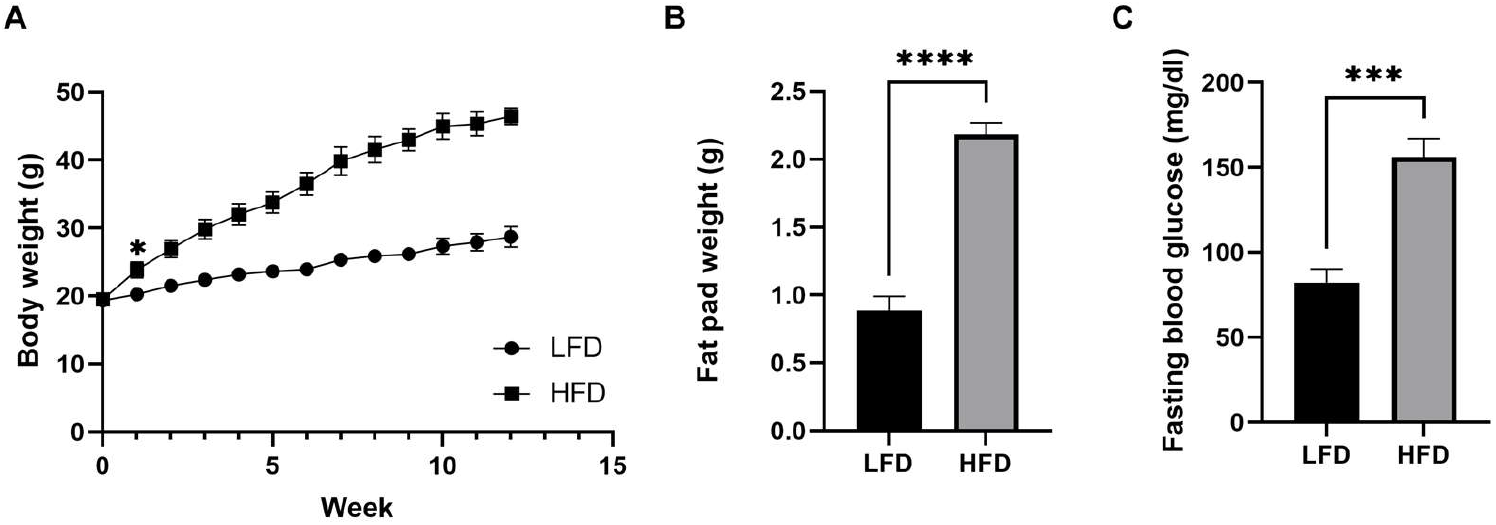
Diet-induced phenotypic variations in male mice. **(A)** Body weight over time. **(B)** Fat pad weight after 12 weeks of diet (n=6/group, mean ± SEM). **(C)** Fasting blood glucose after 12 weeks of diet. Student’s t-test \*\*\**P* < 0.001, \*\*\*\**P* < 0.0001. Data shown as mean ± SEM. n=6 LFD, n=6 HFD.

### 3.2. Liver proteome profile change upon diet-induced obesity

To characterize the holistic changes in the proteome of murine livers during DIO, we performed a global proteomics analysis (Figure 2). In total, 29590 peptides were identified, which could be assigned to 2973 proteins. We then filtered for proteins with LFQ values greater than zero in at least 5 out of 6 replicates in one group. After applying our filtering criteria, 2447 (82%) quantified proteins were retained for further analyses (Supplementary Data Table S1), suggesting high reproducibility and reliability of the proteomic analysis, which was also showcased by distinct clustering in a PCA analysis (Figure 3A). We then performed a student’s t-test to unveil proteins that were significantly regulated after DIO.

**Figure 2.**
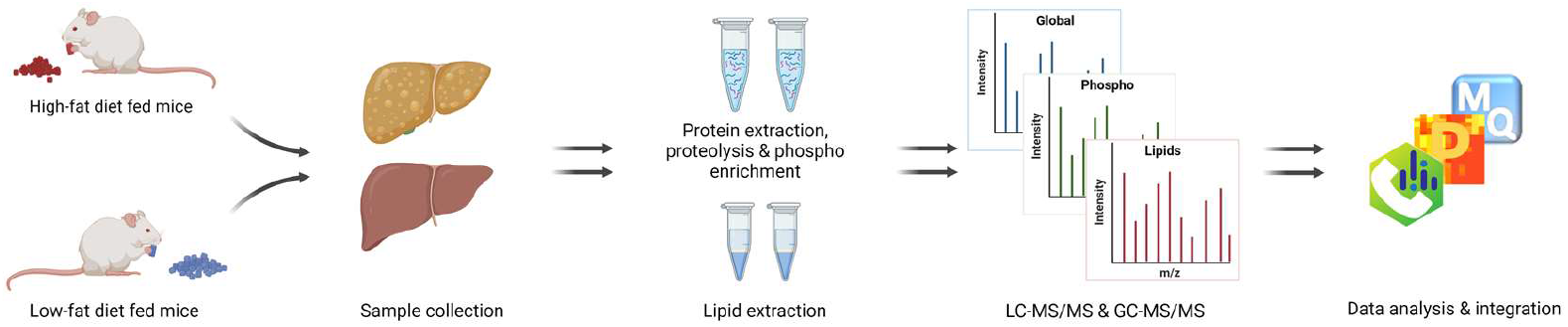
Workflow. Fatty liver establishment by HFD, followed by proteomic, phosphoproteomic, and lipidomic analysis.

**Figure 3.**
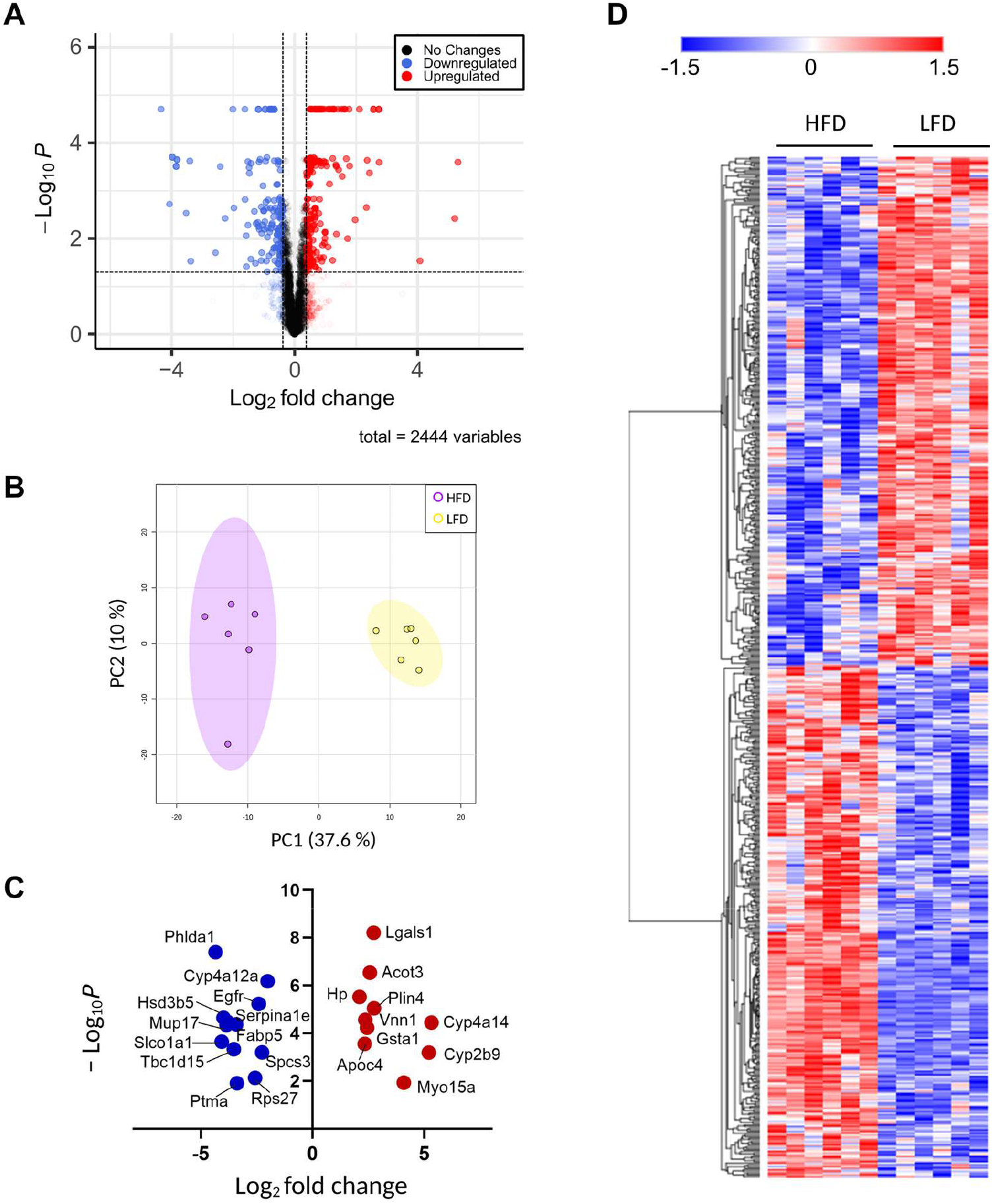
Global proteomic analysis. **(A)** Volcano plot of all quantified proteins. Blue color denotes downregulated proteins, red denotes upregulated proteins, and grey denotes proteins that do not change in response to administered diet. **(B)** PCA plot of each biological replicate. **(C)** Proteins with a t-test q-value less than 0.05 and a |log2Fold-change| greater than 2. Color legend is as described in figure 3B. **(D)** Heatmap representation of all significant proteins.

A volcano plot representation of our filtered data shows that the distribution of upregulated and downregulated proteins was remarkably symmetrical, with 312 proteins above the q< 0.05 and 0.38 log_2_(fold-change) cutoffs, set as the minimum to consider proteins as “significantly regulated” (Figure 3B). Specifically, 141 proteins were downregulated and 171 were upregulated. To gain an insight on the proteins that were most affected by DIO, we separately plotted all significant proteins with an absolute log_2_(fold-change) value greater than 2 (Figure 3C). We observed 11 downregulated proteins and 10 upregulated proteins, many of which are involved in lipid metabolism. Particularly, we observed that Vanin-1 (Vnn1), a GPI-anchored protein that has its expression levels regulated by oxidative stress, and is involved in intestinal inflammation, was among these highly expressed proteins [18,19]. Vnn1 breaks down pantetheine in cysteamine and pantothenic acid, a precursor of coenzyme A. Galectin-1 (Lgals1) was also significantly upregulated in HFD-fed mice, an interesting contrast to our previous results that show Lgals1 is downregulated in adipocytes in response to inflammatory signaling [14]. The clear distinction between the changes in protein levels is further evidenced by a heatmap visualization of the significantly regulated proteins in our dataset, in which two clusters are evident (Fig. 3D).

Thus, to categorize and characterize the proteins significantly regulated during DIO, we performed a Gene Ontology (GO) enrichment analysis of both upregulated and downregulated proteins together. Notably, several GO terms were listed as both downregulated and upregulated, such as monocarboxylic acid and metabolic process, small molecule biosynthetic process, sulfur compound metabolic process and others, which likely indicate a wholistic metabolic reprograming of such processes where specific subsets of proteins are modulated in distinct ways (Figure 4A). For instance, five of the ten proteins with the greatest fold change increase in livers of DIO mice are involved in monocarboxylic acid metabolic process (GO:0032787): Cyp2b9; Cyp4a14; Gsta1; Acot3; and Vnn1 while another protein in the same GO term, Fabp5, is one of the proteins with greatest fold change downregulation (Figure 3C). To easily visualize unique and common pathways in network format, nodes were shown as a pie chart (Figure 4B).

**Figure 4.**
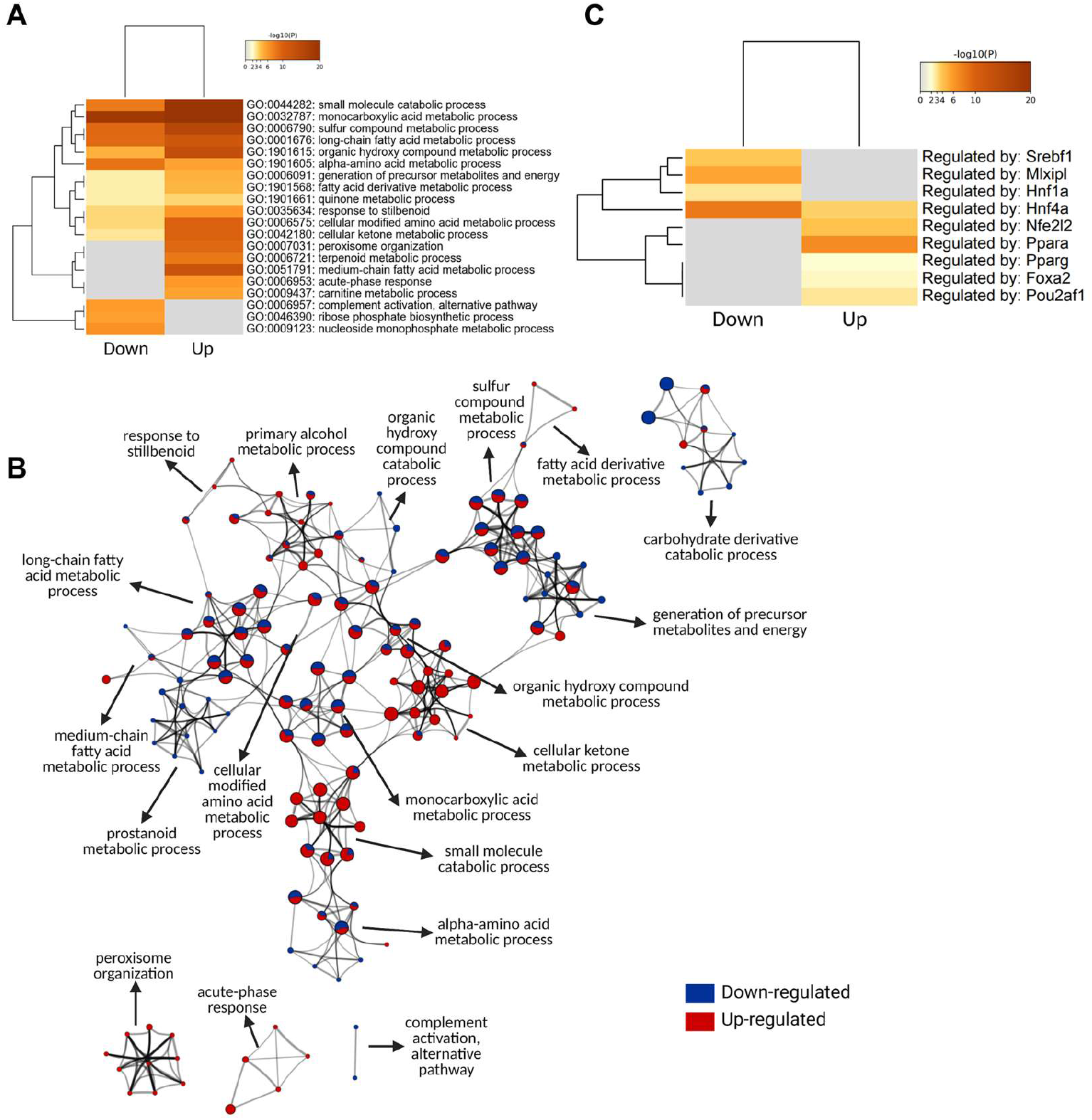
Gene ontology analysis of significantly regulated proteins. **(A)** Metascape enrichment analysis of statistically enriched GO Biological Process (BP) terms. The terms were selected with the best p-value within each cluster as its representative term. **(B)** Enriched clusters GO-BP terms in network format. Each enriched term is displayed as a circle node whose size is proportional to the number of input proteins that fall into the term. Each node is represented as a pie chart and each pie sector is proportional to the number of hits originated from either up-or downregulated proteins. **(C)** Enrichment of transcriptional regulators of significantly regulated proteins using TRRUST (transcriptional regulatory relationships unraveled by sentence-based text-mining) database

Based on TRRUST database, constructed with literature-curated human transcription factor-target interactions [20], we uncovered the most likely transcription factors associated with changes in protein landscape (Figure 3C). The result indicates that Peroxisome proliferator-activated receptor alpha (Pparα) and Nrf2 (Nfe2l2) contribute to DIO-induced protein upregulation while Hnf4α, Chrebp (Mlxipl), Srebp1 (Srebf1) contribute to DIO-induced protein downregulation. The enrichment of the transcription factor Pparα in response to HFD is in line with the observation that Vnn1 is one of the proteins with the highest increase in its protein levels, considering that Vnn1 is a key regulator of Pparα in the liver [19].

### 3.3. Differentially expressed phosphoproteins in the liver from HFD-and LFD fed mice

Protein phosphorylation is a major driver of protein function, protein localization and protein-protein interactions [21,22]. Thus, to gain an insight on how proteins are regulated during DIO in a larger scope, which transcends the changes in protein levels alone, we performed a comprehensive phosphoproteomic analysis of the liver tissues in which the global analysis was performed. We identified a total of 1391 phosphoproteins, containing 1650 class I phosphosites (phosphosites with localization probability > 0.75). We also observed 14 novel phosphosites of Eif3m, Ubxn, Atf7, Reps2, Gprin3, Tacc2, Igf1r, Gsr, Ubxn7, Snx11 and Abcb11 (Supplementary Data Table S2). Of the 1650 class I phosphosites, 1517 (92%) sites were serine, 124 (7.5%) sites were threonine and 9 (0.5%) were tyrosine phosphorylated. Of these 1650 phosphosites, 669 sites were significantly different between HFD and LFD fed mice, only 15 with increased and the remaining with decreased phosphorylation (Supplementary Data Table S2). This result suggests a significant loss of protein phosphorylation in HFD fed mice, highlighting the changes in protein regulatory mechanisms in mice due to DIO. Proteins with increased phosphorylation in HFD included transporter proteins like sodium bicarbonate transporter (Slc4a4), solute carrier anion transporter (Slco1b2), helicase (Rad54l2), RNA binding protein (Rbm39), and glycerol-3-phosphate acyltransferase (Agpat9), etc. Of the 9-tyrosine phosphorylation sites, 6 were significantly different with 1 increased (Slc4a4) and 5 decreased (Gsk3a, Gpxl, Uox, Mapk14 and Arhgap35) phosphorylation. Further, 5 of the 9-tyrosine phosphorylation sites belonged to proteins containing known kinase domains (Mapk1, Mapk14, Gsk3a and 3b, Dyrk1a and 1b and Prpf4b; Supplementary Data Table S2) and phosphorylation of all these kinases decreased in HFD mice liver. We also identified additional 6 phosphoproteins with known kinase domains, and all showed decreased phosphorylation at serine and threonine residues. Predominant decrease in phosphorylation is easily visualized as a heatmap and further evidenced by a volcano plot, representing all filtered phosphosites, in which samples corresponding to HFD are marked by the consistent downregulation of thousands of phosphosites (Figure 5B). Expectedly, the majority of significantly regulated phosphosites identified also showed a similar pattern of decreased phosphorylation of HFD proteins relative to LFD control (Figure 5A).

**Figure 5.**
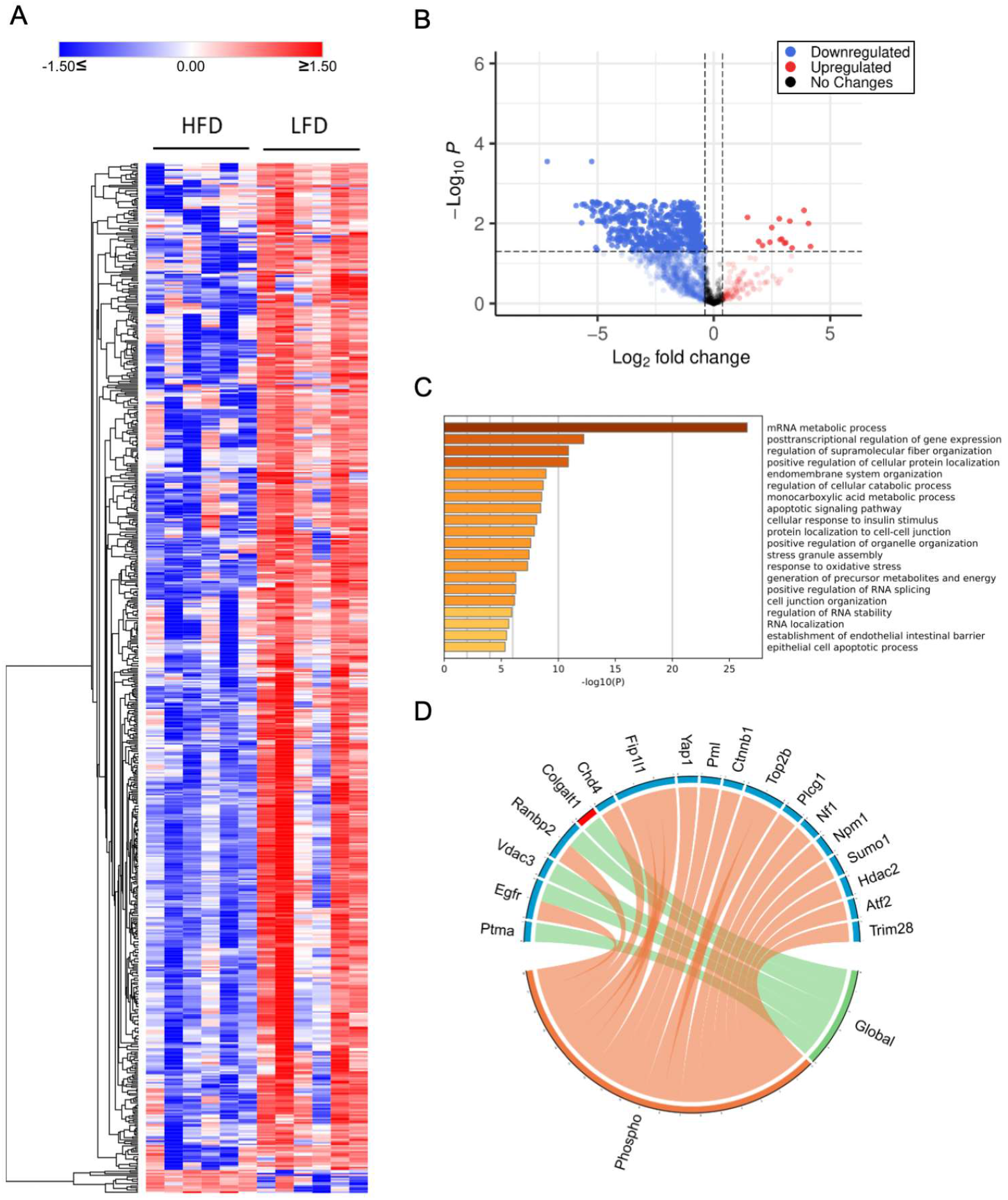
Phosphoproteomic analysis. **(A)** Heatmap representation of all significant phosphosites. Red indicated upregulated phosphosites, and blue indicated downregulate phosphosites. **(B)** Volcano plot of all quantified phosphosites. **(C)** Gene Ontology for biological processes enriched from proteins with significantly downregulated sites. **(D)** PML proteins significantly regulated at phospho and global levels

Among the many downregulated phosphoproteins in HFD mice liver were several solute career (Slc) transporters including Slc2a2, Slc10a1, Slc16a1, Slc16a10, Slc16a7, Slc33a1, Slc38a4, Slc4a1, Slc4a4, and Slc4a10 (Supplementary Data Table S2). These transporters facilitate the transport of substrates across cell membranes, including glucose, inorganic and organic ions, small molecule drugs, xenobiotics and amino acids, and contribute to insulin signaling, glucose homeostasis and the etiology of different metabolic diseases [23,24]. The glucose transporter isoform Glut2 (known as Slc2a2) was phosphorylated at serine 522, and phosphorylation decreased in HFD compared to LFD mice liver (Supplementary Data Table S2). Glut2 expression was also down at the protein level, suggesting impaired glucose homeostasis and dysregulated insulin response. Other notable downregulated solute transporters were lactate and H^+^ transporter, Slc16a1; acetyl-CoA transporter, Slc38a1; sodium and amino acid transporter, Slc33a1; and sodium and bile acid transporter, Slc10a1. We also identified downregulation of acetyl-CoA carboxylase (Acaca and Acacb), both at the protein and phosphoprotein levels (Supplementary Data Table S2). Acaca was phosphorylated at serine 23, 29 and 79, whereas Acacb was phosphorylated as serine 1332. The decreased phosphorylation of the enzymes in all these phosphosites under HFD fed condition further indicates dysregulation of lipogenesis and glucose homeostasis.

Gene Ontology (GO) enrichment analysis of differentially expressed phosphoproteins indicated that RNA metabolism was highly affected by HFD at the phosphorylation level. Interestingly, we also found that cellular response to insulin stimulus was one of the top 20 most significant downrelgulated processes among the enriched phosphoproteins (Figure 5C). Many other biological processes, as expected from previous studies on diet-induced obesity [8,25,26], are related to cellular metabolic and catabolic processes, membrane and organelle organization, cell-cell communication, protein localization, apoptotic signaling, and oxidative stress responses (Fig. 5B).

Recent studies have reported that the tumor suppressor promylocytic leukemia (Pml) protein plays a regulatory role in cellular metabolism, by controlling PPAR, which, in turn, is involved in an important signaling pathway that modulates lipid homeostasis [27,28]. PML is known as the key organizer of the PML-Nuclear Bodies (PML-NBs), and has fascinated scientists for many years due to its multifaceted role under many cellular conditions, notably acute promyelocytic leukemia [29,30]. Liver Pml ablation induces extensive reprograming of metabolic pathways, including an accelerated fatty acid metabolic rate accompanied by decreased total lipid accumulation in the liver, as well as insulin resistance [31]. To investigate the effects of HFD in PML-NB proteins, we constructed a library using protein-protein interaction databases, and filtered for proteins identified in our global and phosphoproteomics study. Many well-known PML-NB proteins were identified (Fig 5C), including the protein Pml. Phosphorylation of Pml was downregulated, notably at S17, a phosphosite targeted by the kinase Cdk. (Supplementary Data Table S2). Further, several E3 sumo and ubiquitin ligases were also identified as phosphoproteins and were downregulated in the HFD mice liver compared to the LFD mice liver, including RanBP2, Nedd4, Praja-1, Rbbp6, Urb4, and Zfp19 (Figure 5D). These results suggest that not only protein phosphorylation, but proteins may be regulated by sumoylation and ubiquitination.

### 3.5. Differentially expressed lipids in the liver from HFD-and LFD fed mice

Proteome and phosphoproteome analysis alone might not reflect the exact changes in lipid content of the liver. To directly investigate the changes in lipid composition and regulation at the metabolite level due to HFD, we conducted an untargeted global lipidomics using LC-MS and GC-MS analytical systems. Untargeted lipid profiling detected a total of 3728 lipid ions present in at least 80% of one group. Data processing using MS-Dial allowed for the tentative identification of 1801 features after blank filter. 464 lipid ions had MS2 acquired data and were used for statistical analysis (Supplementary Data Table S3). Most lipids detected were triacylglycerols (TAGs) and phosphatidylcholines (PCs), followed by diacylglycerols (DAGs) and sphingolipids. Several lipids were suggested with Riken IDs and grouped as unknowns. FAMEs (here on called FFA) were identified using GC-MS and normalized to C17 internal standard (Supplementary Data Table S4). Lipid data was centered at the mean and divided by the standard deviation of each variable to scale it and account for the difference in intensities in the subsequent analysis.

Unsupervised PCA analysis of the lipid features relative intensities revealed clear separation of the two groups with a 42.4% of explained variance (Fig 6A). The heat map illustrates the difference in the detected lipid profile of each sample and shows two main clusters with opposite trend (Fig 6B). Univariate analysis comparing lipid profiles of HFD and LFD fed mice identified 109 lipid features with p-values below 0.05 after t-test and 42 features with reduced and 67 with increased relative amounts of at least 2-fold-change, which are represented in the volcano plot (Fig 6C). As expected, TAGs and FFA were increased in HFD compared to LFD and most PCs, PE and LPCs were decreased. Lipid ontology (LION) enrichment [32] was performed for lipid class overrepresentation analysis (Fig 6D), and showed upregulated TAGs, followed by glycerolipids and ether glycerophosphocholines. On the other hand, glycerophospholipids, including lysoglycerophospholipids and ceramides. Although not overrepresented, unusual acylcarnitines CAR(24:6) and CAR(28:6) were significantly downregulated. Glycerophosphoglycerols, such as BMP (35:4) and HBMP (48:0) were significantly reduced in HFD compared to LFD despite not being a largely represented. LION analysis for cellular component, function, and physical and chemical properties (Fig 6E), showed that lipid storage and lipid droplet formation associated lipids were significantly increased in DIO mice as well as lipids with neutrally charged head group. Also, 12 to 22 carbon fatty acids either saturated, monounsaturated or with more than 3 double bonds were increased, while 24 to 26 carbon fatty acids with 2 and 3 double bonds were reduced compared to LFD. Lipids linked to endoplasmic reticulum (ER) and lipid-mediated signaling were significantly reduced in HFD compared to LFD, as well as lipids associated to the membrane components, intrinsic curvature and positive/zwitterion head group.

**Figure 6.**
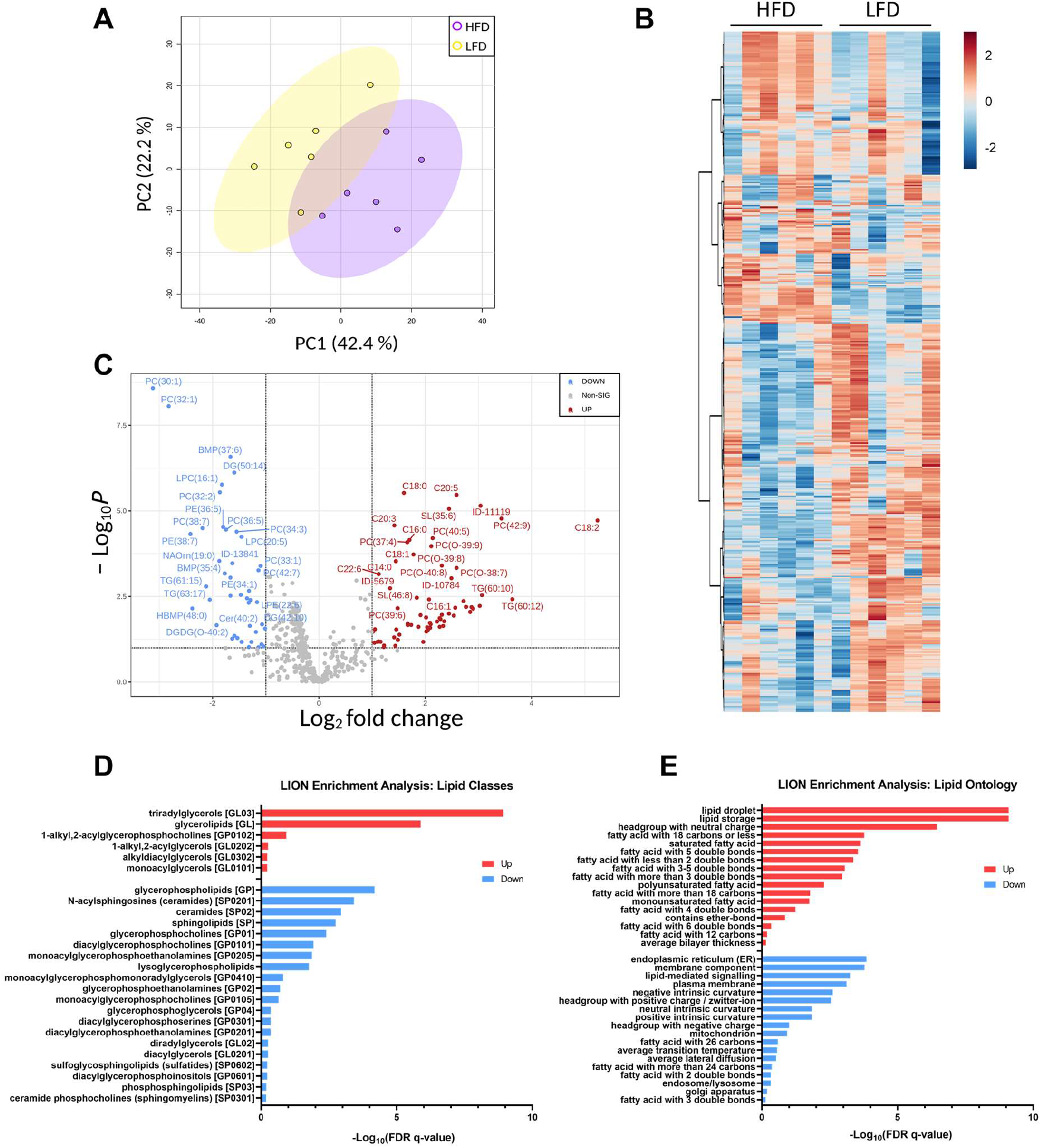
Lipid profiles. (**A**) PCA plot of each biological replicate. (**B**) Heatmap representation of all significantly changing lipids. Red indicated upregulated lipid species, and blue indicated downregulated lipid species. (**C**) Volcano plot of all quantified lipids. (**D**) Principal lipid classes changed in the two groups. (**E**) Lipid ontology enrichment for function, cellular compartment and chemical and physical properties significantly regulated.

To derive insights on whether distinct lipid components align with the changes in proteins or phosphoprotein profiles, correlations between detected lipid species and significantly changing proteins in the global and phophosproteomics were analyzed. Lipids were assigned to their respective classes, and the class correlation was obtained as described by Chauhan *et al*. [33]. Variable importance in projection (VIP), which is defined as the weighted sum of squares of the loadings in a partial least squares discriminant analysis that takes into account the amount of explained variation in each dimension, was used to rank the features. Correlation heatmaps of global proteins lipid classes showed that glycerophosphoglycerols, ether-linked digalactosyldiacylglycerols, LPC, LPE, neutral sphingolipids and PS presented stronger correlation (p<0.0 and VIP >0.8) with the identified proteins, while triacylglycerols did not appear highly correlated with global proteins ((Fig 7A; Supplementary Data Table S5). When examining the phosphoproteomics correlation heatmap (Fig 7B; Supplementary Data Table S6), fewer lipid classes had abundant significant correlation across the phosphosites identified. Ether-linked PC (EPC) and FFA were predominantly negatively correlated with phosphosites such as Tnks1bp1 (S796), Chd4 (S508), Grb7 (S420), Acss2 (S263), Bad (S155) while acylcarnitines, glycerophosphoglycerols, LPC and LPE were mostly positively correlated with the exception of Thrap3 (S669), Agpat9 (S68), Hadh (S13), Pdha1 (S300), Prpf4b (S145), Rad54l2 (S1168 and S1171), Rbm39 (S127 and S129), Rpap3 (S429), Scaf4 (S154), Slc4a4 (S68), Slco1b2 (S290), Sord (S169), Srsf2 (S189 and S191), and Tnks1bp1 (S866) (Fig 7B). To gain insights on potential biological implications of strong correlation between certain lipid classes and phosphoproteins, significantly different proteins with a VIP larger than 1 that are in significant correlation with lipids were analyzed using REACTOME for pathway analysis including interactors. The identified significant pathways were comprised in the main categories of metabolism, programmed cell death, signal transduction, transport of small molecules, disease, and metabolism of proteins. Different pathways involved in energy metabolism were highlighted such as glycolysis, gluconeogenesis, ChREBP activates metabolic gene exression, and PKA-mediated phosphorylation of key metabolic factors, as well as PP2A-mediated dephosphorylation of key metabolic factors (Fig 8A). Pathways related to mitochondria function including activation of PPARGC1A (PGC-1alpha) by phosphorylation and carnitine metabolism were also significantly enriched (Fig 8A). Several proteins were grouped under translation by ribosomal scanning and start codon recognition, and signaling by Rho GTPases (Fig 8A). Interestingly, when individual phosphorylation sites classified under certain pathway were categorized by correlation coefficient with each lipid class (Fig 8B), it was evident that not all the phosphosites had the same behavior. For example, Eif3b had 4 different phosphosites that meet the correlation cut-off criteria but each one of those sites showed different correlations with AC, FFA, Ether-PC, LPC and LPE, highlighting specificity of phosphosites and their unique regulation within a protein. Another similar example is a membrane protein Slc4a4. Slc4a4 (S68) had negative correlation with phospholipids but such relationship was not observed with Slc4a4 (Y64).

**Figure 7.**
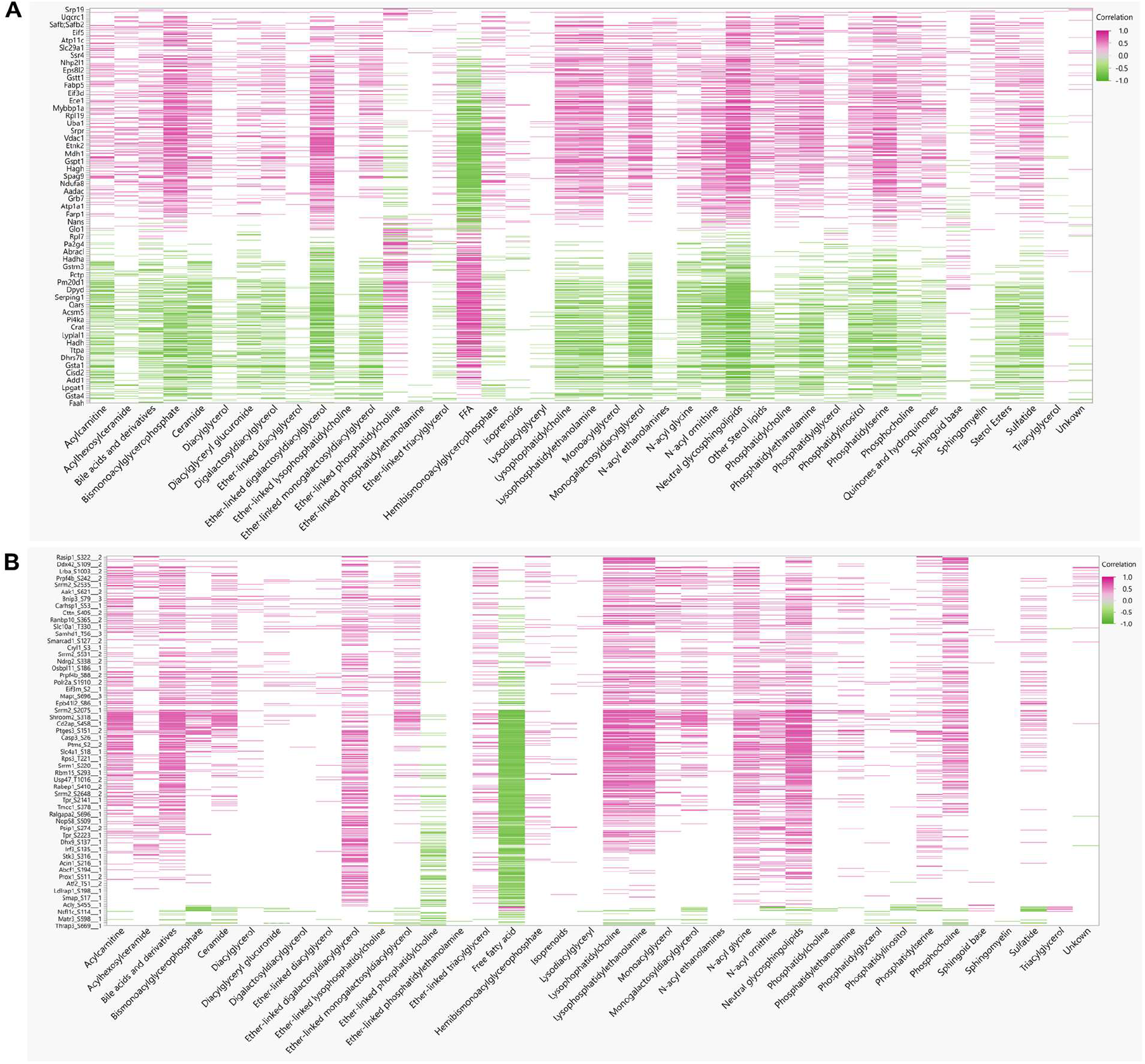
Protein:Lipid correlations. (**A**) Pearson correlation matrix between 38 lipid classes and 2447 proteins or (**B**) 649 phosphoproteins.

**Figure 8.**
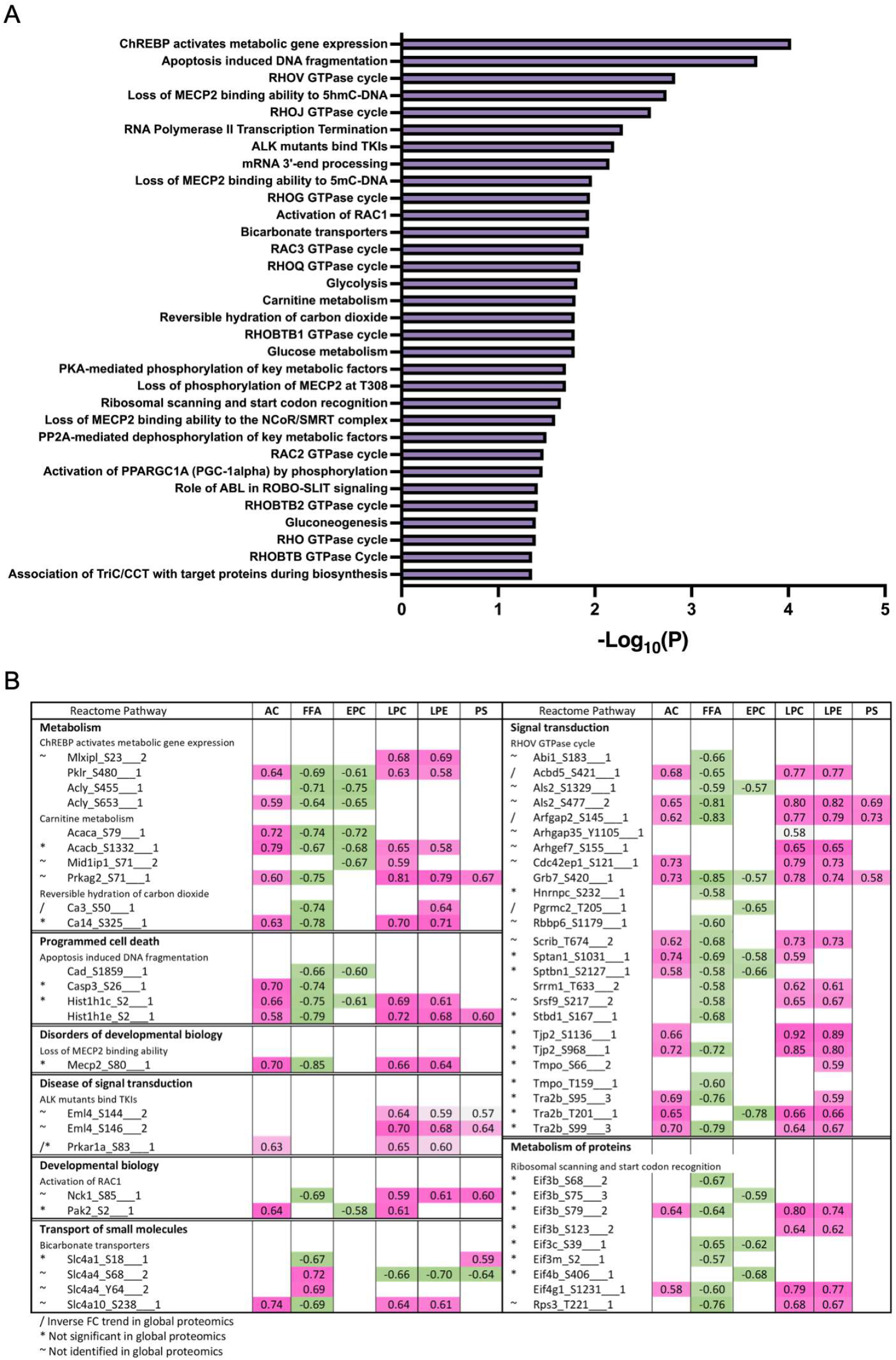
Reactome enrichment of correlation analysis for phospho datasets. (**A**) Reactome enriched pathways from global phosphoproteins significantly correlated with lipids. (**B**) Correlation analysis of phosphosites of interest, categorized by enriched reactome pathway, and lipid classes.

## 4. Discussion

Applying multi-omics approach, we identified significant modulation of the liver proteome and phosphoproteome involved in signaling implicated in obesity and insulin resistance. We observed loss of phosphorylation in the majority of identified proteins in HFD-mice, suggesting that metabolic disorders due to HFD induced obesity are characterized by dysregulated phosphorylation-dependent signaling. Further, decreased phosphorylation of many protein kinases and key metabolic enzymes involved in lipid and glucose homeostasis imply underlying functional consequences of obesity and insulin resistance.

### 4.1. Altered proteomes related to lipoprotein assembly, lipoprotein uptake, de novo lipogenesis (DNL) and fatty acid uptake

Liver predominantly produces lipoproteins, other than chylomicron. DIO animals had abundant apolipoproteins (ApoA4, ApoC1, ApoC2, ApoC4, ApoE, ApoH), which suggests an increased level of circulating lipids. In terms of cellular uptake of blood lipids, fatty acid uptake protein Cd36 was highly abundant in livers of obese animals. A previous study showed Cd36 null mice are protected from hepatic steatosis induced by Lxr agonists and that Cd36 is a common target of Lxr, Pparγ and pregnane X receptor (Pxr) [34]. While we did not identify Pparγ, we identified up-reguation of Pparα, which belongs together with Pparγ as a nuclear receptor subfamily, and may contribute to Cd36 upregulation in DIO animals (Fig 4C).

Three key rate-limiting enzymes in de novo lipogenesis (DNL) (Acly, Acaca, Fasn) as well as Scd1 that converts saturated fatty acids into monounsaturated fatty acids in the final step of DNL were present in lower amount in the livers of obese mice. Acly was also found to be dephosphorylated at Ser455 in obese animals, adding another layer of DNL attenuation as phosphorylation at Ser455 is known to increase Acly enzymatic activity [35]. On the other hand, despite Acaca protein level decreased (Log2FC: −0.83677), loss of phosphorylation at regulatory site Ser79 was observed at a greater extent (Log2FC: −1.289), suggesting Acaca is enzymatically in an active state although protein level may be less in livers of DIO because phosphorylation is known to inhibit Acaca activity [36].

Other lipogenic enzymes such as desaturases that introduce double bonds in acyl chains (Fads1, Fads2) and elongases which elongate C16 to longer acyl chain (Elovl2, Elovl5) were identified but did not show a difference between lean and obese animals. Meanwhile, lipid droplet proteins (Plin2, Plin4) were present in higher amount in obese animals, indirectly showing heightened lipid storage. Hepatic DNL is mainly controlled by transcriptional regulation of genes by Srebp1 [37], Chrebp [38], liver X receptor α [39] and Pparγ [40]. In agreement with the finding, Srebp1 and Chrebp were shown as enriched transcription factors responsible for downregulated proteins according to TRRUST analysis (Fig 4C). Overall, lipid accumulation in livers of HFD-fed animals is largely attributed to increased lipid flux rather than DNL.

Impaired glucose balance and insulin resistance characterize metabolic disorders in DIO. There are growing evidences that Slc transporters contribute to the etiology of various metabolic diseases [41]. Slc transporters are located in membranes and are highly expressed in the liver, kidney, heart, gut and brain, and are emerging as potential drug targets [24]. They serve as ‘metabolic gate’ of cells and mediate the transport of a wide variety of nutrients and metabolites such as glucose, amino acids, vitamins, neurotransmitters, inorganic ions, organic anions, metals, and amino acids. Among the 35 identified Slc transporters, expression of 12 members showed significant changes between HFD and LFD fed mice liver, of which 2 members (Slc22a1 and Slc25a10) had increased expression and 10 (Slc10a1, Slc25a11, Slc25a20, Slc25a22, Slc25a3, Slc2a2, Slc33a1, Slc6a13, Slc9a3r1 and Slc01a1) had decreased expression. We identified several of these Slc transporters with loss of phosphorylation suggestive of defects in insulin signaling and transport of glucose and other molecules to membranes, essential for normal cellular function. The transporter Slc2a2 is responsible for the transport of glucose into β-cell and facilitates glucose-mediated insulin secretion and signaling. Recessive mutation of Glut2 gene in mouse showed hyperglycemia and abnormal glucose homeostasis [42]. Its downregulation and loss of phosphorylation in our study further suggest impaired utilization of glucose. However, the functions of other identified Slc transporters and the consequences of their loss of phosphorylation in obesity are unknown.

### 4.2. Increased mitochondrial fatty acid β-oxidation, ketone body formation

Intracellular fatty acids are activated by acyl-CoA synthetase before being channeled into different metabolic fates such as β-oxidation, triacylglycerol synthesis or phospholipid synthesis. Our results show Acsl1, predominant isoform in the liver, was present more in obese animals but less than our cut-off value (log2 FC: 0.306). In sharp contrast, Acsl4 was significantly less abundant in obese animals (log2FC: −1.478). Previous study showed arachidonic acid is a preferred substrate for Acsl4 [43] and that Acsl4 protein expression is controlled by substrate induced post-translational regulation in which arachidonic acid promotes ubiquitin-proteasomal degradation [44]. Our fatty acid measurement (Supplementary Data Table S3) shows a high level of arachidonic acid in livers of DIO animals, and this may have contributed in part to reduced abundance of the protein. Other acyl-CoA synthetases with significant fold-change include Acss3 (log2FC: 1.64) and Acss2 (log2FC: −0.97). Acss3 is a mitochondrial enzyme whose expression is upregulated under ketogenic conditions [45] and our data shows abundance in livers of DIO animals. On the other hand, Acss2 was not only less abundant but also showed lower phosphorylation levels (Ser30, Ser263, Ser267) in livers of obese mice. Suppression of Acss2 in the liver after HFD feeding was reported previously [46] and others reported mice lacking Acss2 are protected from steatosis induced by DIO [47], suggesting downregulation of Acss2 may serve as a defense mechanism against excess fat storage in liver. However, it is unlcear how phosphorylation affects the enzymatic activity and future study is warranted.

Our results are consistent with previous reports on the proteomic landscape of hepatic steatosis by showing an abundance in proteins involved in mitochondrial β-oxidation. Carnitine shuttling enzymes (Cpt1α, Slc25a20) were more abundant in livers of obese animals, transporting excess fatty acids from the cytoplasm into the mitochondrial matrix. Acadm and Acad11 enzymes that catalyze dehydrogenation of fatty acyl-CoAs to form enoyl-CoA were also more abundant in obese animals, while the levels of Acadl were not significantly different between the two groups. Acads and Acadvl were present at a higher level in the livers of obese animals but did not meet our fold change cut-off value. The enzymes that catalyze the hydration of the enoyl-CoAs (Ech1, Hadh) are abundant in obese animals, with upregulation of Hadh Ser13 phosphorylation.

Hepatic ketogenesis occurs in the mitochondria and is activated to convert excess acetyl-CoA generated from β-oxidation into ketone body intermediates. Several proteins involved in ketogenesis were upregulated in the liver of HFD fed animals: Bdh1; Acat1; Hmgcs1,2; and Hmgcl. Aacs that converts ketone bodies into acetoacetyl CoA for cholesterol or fatty acid synthesis were less abundant in HFD fed animals.

### 4.3. Increased abundance of proteins involved in peroxisomal β-oxidation and microsomal ω - oxidation

Peroxisomes are single membrane-enclosed subcellular organelles, particularly abundant in hepatocytes. One dynamic metabolic process that peroxisomes participate in is the degradation of fatty acids that cannot occur in the mitochondria, in specific (1) very-long chain fatty acids with at least 22 carbons; (2) branched-chain fatty acids; (3) bile acid intermediates such as di- and trihydroxycholestanoic acids; (4) long-chain dicarboxylic acids [48]. Peroxisomal acyl-CoA oxidases that participate in catalyzing peroxisomal β-oxidation (Acox1, Acox2, Acaa1a, Acaa1b), as well as the bifunctional protein Ehhadh that catalyze the hydration and dehydrogenation step of β-oxidation, were all upregulated in obese animals. Acyl-CoAs that are produced during peroxisomal β-oxidation can be hydrolyzed into fatty acids by Acot4, or, alternatively, converted to carnitine esters and free CoA by Crat and Crot, all of which were more abundant in HFD-fed mice. Accordingly, our results show that proteins involved in peroxisome biogenesis were abundant in obese mice, suggesting increased demand for the maintenance and biogenesis of the organelle to support increased peroxisomal function. Peroxisome organization (GO:0007031) was enriched uniquely in livers of DIO animals (Fig 4A) and especially, proteins involved in peroxisome biogenesis known as peroxins (Pex1, Pex11a, Pex16, Pex19, Pex3, Pex6) were present more in obese animals.

Another lipid catabolism pathway that increased in response to elevated hepatic lipid overflow was microsomal ω-oxidation. The fatty acid ω-hydroxylase Cyp4a10 and Cyp4a14 were highly upregulated in obese animals (Fig 3C). The resultant fatty ω-hydroxy acids are catalyzed by two subsequent reactions by alcohol dehydrogenase and aldehyde dehydrogenase. Our results show that microsomal fatty aldehyde dehydrogenase Aldh3a2 was also more abundant in obese animals. Overall, HFD feeding stimulated mitochondrial, peroxisomal and microsomal fatty acid oxidation systems. The fact that the key transcriptional factor Pparα is most enriched in DIO animals according to TRRUST analysis (Fig 4C) further supports upregulation of β-oxidation in three different subcellular organelles.

### 4.4. Decreased glycolysis and increased gluconeogenesis

5’-AMP-activated protein kinase (AMPK) is an energy sensor kinase that activates energy-producing pathways when the intracellular ATP level is low. In the livers of obese mice, AMPK catalytic alpha-1 subunit, Prkaa1 (S496) had reduced phosphorylation, as well as non-catalytic gamma subunit Prkag2 (S71, S87, S90), Prkab1(S108, S96), indicating reduced kinase activity. Prkag1 was also less abundant in livers of DIO. Accordingly, we identified that several proteins under AMPK regulation showed reduced phosphorylation levels in obese animals. For example, glycogen synthase 2 (Gys2) phosphorylation at Ser8 (log2FC: −1.7954) and Ser 627 (log2FC: −4.591) was downregulated in DIO, indicating the active state of the enzyme, favoring glycogen synthesis despite decreased protein level (log2FC:-0.7757).

Meanwhile, decreased phosphorylation states of glycolytic enzymes that are under the control of AMPK induce glycolysis inhibition. Our data shows downregulated phosphorylation of Pfkfb1 at Ser33, which indicates the activation of bisphosphatase and degradation of fructose-2,6-biphosphate, favoring gluconeogenesis [49]. Also, Pfkl reduced phosphorylation at Ser775 proteins further supports increased gluconeogenesis by decreasing kinase activity [50]. Carbohydrate response element-binding protein (Chrebp), which regulates gene expressions involved in glycolysis, gluconeogenesis and DNL, is one of the main transcription factors enriched in our downregulated protein dataset (Fig 3C). Consistently, our phosphoproteome result shows Chrebp phosphorylation was decreased at Ser514 in DIO, the residue critical for maintaining the transcriptional activity of Chrebp by enhancing Chrebp O-GlcNAcylation via phosphorylation [51]. Accordingly, one of the Chrebp target gene, Pklr, was less abundant in DIO animals, supporting decreased glycolysis.

### 4.5. Altered abundance of proteins involved in ROS scavenging

The influx of free fatty acids into hepatocytes leads to increased ROS, such as hydrogen peroxide (H_2_O_2_) and superoxide anion (O2_•_−), potentially increasing impairing intracellular oxidative stress. In fact, NAFLD pathogenesis is explained by a double hit model, in which the first hit is steatosis, sensitizing the liver to the second hit, in the form of ROS and cytokines [52]. Mice lacking Nrf2, a key transcriptional factor that controls the antioxidant system, were prone to NASH when challenged with methionine- and choline-deficient diet [53]. Our proteome results are consistent with previous studies which show the induction of antioxidant systems in the steatotic liver. Glutathione peroxidase 4 (Gpx4) is an enzyme found mostly in the mitochondria, which directly detoxifies membrane lipid peroxides at the expense of glutathione. Gpx4 was more abundant in the livers of HFD-fed mice. Glutathione synthetase (Gss), that replenishes glutathione from glycine and γ-glutamylcysteine, was also upregulated in DIO. Furthermore, in our TRRUST analysis, Nrf2 was upregulated (Fig 4C). In DIO, the glutathione-S transferases (Gst) targets of Nrf2, including Gstm1,2,3,4, Gstk1, Gsta1,4 and Gstt3, were more abundant. Another Nrf2 target, Nqo1 was also more abundant in DIO animals. However, few Gst isoforms were less abundant in DIO: Gstm7; Gstp1,2; and Gstt2, making cellular modified amino acid metabolic pathway (GO:0006575) a common GO term in both lean and DIO groups. A future investigation needs to identify the biological significance of differential induction of multiple Gst isoforms during DIO.

### 4.5. Regulation of PML-NB proteins under HFD

While the role of Pml as a tumor suppressor protein is well known, its role in cellular metabolism, notably under obesity or HFD conditions, is unknown or inconclusive. Carracedo and Pandolf [54] reported increased hepatic Pml protein levels during liver steatosis. The Pml protein interferes with liver metabolism and controls fatty acid oxidation in stem cells [55], and the depletion of Pml in mice decreases liver fatty acid accumulation after a long-term western diet [31]. In contrast, other studies showed that Pml inhibits adipogenesis, and loss of Pml results in fat accumulation in mice [27]. It is possible that differences in a mouse strain, diets, aging, and environment may contribute to the inconsistent phenotypes [31,56]. Our results also emphasize the role of Pml in diet-induced obesity and cellular metabolism, and its regulation by phosphorylation. This PTM-dependent regulation may explain some of the inconsistencies reported in previous studies as none of those studies investigated the hepatic function of phosphorylated Pml. Downregulation of many E3 sumo and ubiquitin ligases also suggest the importance of these modifications, and further highlight the regulation of Pml and other NB proteins via diverse modifications as phosphorylation, SUMOylation and ubiquitination are known to be interdependent to each other to execute cellular functions under various stressed conditions [57,58].

## 5. Conclusions

Our integrated proteomic and lipidomic study of HFD induced fatty liver samples from animal models identified a spectrum of biological pathways that were significantly altered. Using this holistic multi-omic approach, we showed how changes in lipid composition and quantity are correlated with the changes in the liver proteome and phosphoproteome or vice-versa. In particular, we showed correlation between site specific phosphorylation changes with the changes in lipid classes. These results offer new opportunities for improving our understanding of the shift in liver metabolism of mice fed excess calories. This study not only underscores the mechanistic insights about altered liver metabolism and health risk associated with HFD, but also provides data that could be useful for future investigations that aim to find new treatment to control the onset or slow down the progression of HFD induced hepatosteatosis.

## Abbreviations for lipid classes

TAG: triacylglycerol
DAG: diacylglycerol
AC: acylcarnatine
FFA: free fatty acid
FAME: fatty acid methyl ester
PC: phosphatidylcholine
PE: phosphatidylethanolamine
PS: phosphatidylserine
LPC: lysophosphatidylcholine
LPE: lysophosphatidylethanolamine
EPC: ether-linked phosphatidylcholine

## Supplementary Materials

The following supplementary data are available online at www.mdpi.com/xxx/s1, Table S1: List of all proteins identified in global proteomic analysis.. Table S2: List of all phosphoproteins and phosphosites identified in phosphoproteomics analysis. New phosphosites are indicated by blank in “known site” column. Table S3: List of all lipid features identified. Table S4: Corrrelation analysis between total proteins and lipids. Table S5: Correlation analysis between phosphoproteins and lipids. Table S6: Normalized relative abundance of different lipid classes identified. Table S7: List of all identified free fatty acids in different samples with different carbon chain length and relative expression in different samples.

## Author Contributions

Conceptualization, methodology, sample preparation and data analysis, S.Q.K, R.M., J.F. and U.K.A.; animals, K.K.B., other resources, U.K.A., original draft preparation, S.Q.K., R.M., J.F. and U.K.A.; writing review and editing, K.K.B., K.H.K. and U.K.A.; supervision, U.K.A.; project administration, U.K.A.; funding acquisition, U.K.A. All authors have read and agreed to the published version of the manuscript. S.Q.K., R.M. and J.F. contributed equally to this work.

## Institutional Review Board Statement

The study was approved by Purdue University’s IACUC protocol (protocol number1111000154, approved 12/7/2008-12/06/2024).

## Data Availability Statement

Raw LC-MS/MS data files can be accessed through MassIVE (https://massive.ucsd.edu/) with the ID: MSV000088835.

## Acknowledgments

All the LC-MS/MS data were collected at the Purdue Proteomics Facility, Bindley Bioscience Center. S.Q.K. received graduate research assistantship from the Bindley Bioscience Center, Purdue University in 2020-2021. R.M. was supported through funding from the Showalter Trust Foundation. We thank Theresa D’Aquilla for assistance with mice care and liver collection. Illustrations were created with BioRender.com

